# Optogenetics-integrated gut organ culture system connects enteric neurons dynamics and gut homeostasis

**DOI:** 10.1101/2024.03.28.587149

**Authors:** Gitali Naim, Hadar Romano, Sivan Amidror, David Jessula Levy, Adva Cohen, Carmel Sochen, Yasmin Yarden, Mengyang Feng, Nairouz Farah, Rotem Tsentsarevsky, Ziv Brodie, Yasmin Reich, Ariel Simon, Einat Toister, Irit Shoval, Yossi Mandel, Moshe Biton, Nissan Yissachar

## Abstract

The enteric nervous system (ENS) senses microbiota-derived signals and orchestrates mucosal immunity and epithelial barrier functions, in health and disease. However, mechanistic dissections of intestinal neuro-immune-microbiota communications remain challenging and existing research methods limit experimental controllability and throughput. Here, we present a novel optogenetics-integrated gut organ culture system that enables real-time, whole-tissue stimulation of specific ENS lineages, allowing for detailed analysis of their functional impact. We demonstrate that optogenetic activation of enteric cholinergic neurons rapidly modulates intestinal physiology. Interestingly, distinct neuronal firing patterns differentially modulate neuro-immunological gene expression and epithelial barrier integrity. Furthermore, diverse enteric neuronal lineages exert distinct regulatory roles. While cholinergic activation promotes gene-sets associated with type-2 immunity, tachykininergic enteric neurons differentially control mucosal defense programs. Remarkably, luminal introduction of the immunomodulatory bacterium *C. ramosum* significantly remodeled cholinergic-induced neuro-immunological transcription. These findings suggest that complex combinatorial signals delivered by gut microbes and enteric neurons are locally integrated to fine-tune intestinal immunity and barrier defense. Collectively, we provide a powerful platform for systematic discovery and mechanistic exploration of functional neuroimmune connections, and their potential modulation by drugs, microbes, or metabolites.

**Short abstract:** The enteric nervous system senses microbiota-derived signals and orchestrates mucosal immunity and epithelial barrier functions. Mechanistic dissections of intestinal neuro-immune-microbiota communications remain challenging. We developed an optogenetics-integrated gut organ culture system for real-time neuronal stimulation and analysis. We revealed neuronal-specific activity patterns, which differentially regulate intestinal transcription and epithelial barrier integrity. Collectively, we provide a powerful platform to test neuroimmune connections and their potential modulation by drugs, microbes, or metabolites.

## Main text

Homeostatic functions of organs and tissues critically rely on multidirectional and dynamic interactions between diverse cellular systems. However, mechanistic dissections of intercellular connections are often limited by methodological constraints. *In vitro* cultures cannot faithfully recapitulate the cellular complexity and spatial organization observed in tissues, and *in vivo* animal models are limited by experimental readouts. This is especially evident in the gut, where intricate communications between neuronal, immunological, and epithelial compartments with the gut microbiota, are integrated to maintain intestinal functions in steady state^1,2^. For example, enteric neurons sense microbial signals and subsequently modulate innate lymphoid cells (ILC) ^3–6^, macrophages and monocytes^7–9^, effector and regulatory T cells^10,11^, and the epithelial barrier^12,13^, collectively orchestrating barrier defense and mucosal immunity. Thus, there is an unmet need for an experimental platform that facilitates whole-tissue, multi-parametric and real-time perturbations of enteric nervous system (ENS) activity and luminal microbiota content, to promote the identification and mechanistic dissections of novel intercellular communications modules.

To address this gap, we devised an optogenetics-integrated gut organ culture system for precise stimulation of specific enteric neuronal lineages in multiplexed culture of whole intestinal fragments (**Fig.1A**). For this purpose, we modified the gut organ culture device we previously developed ^10,14^. The new device is opaque, to prevent light transfer between multiple gut fragments cultured in individual channels and is sealed by a 3D-printed lid that incorporates LED light diodes and O2/CO2 input ports (**Fig.1A**; **FigS1A-B**). LED light/dark cycles, frequency and duration in each channel are controlled by an Arduino open-source electronic platform (**Fig.S1C-D**) and custom-made user interface software developed in-house. The gut culture device supports parallel culture of 6 intestinal fragments, each connected to input and output ports. Computer-controlled syringe pumps maintain luminal flow (to facilitate the coordinated introduction of microbes or compounds) and continuously replenish the extra-intestinal culture medium (to support cultured tissues viability). We have previously demonstrated that these culture conditions maintain tissue viability, structure, and cellular components for up to 24h^10^.

**Figure 1:**
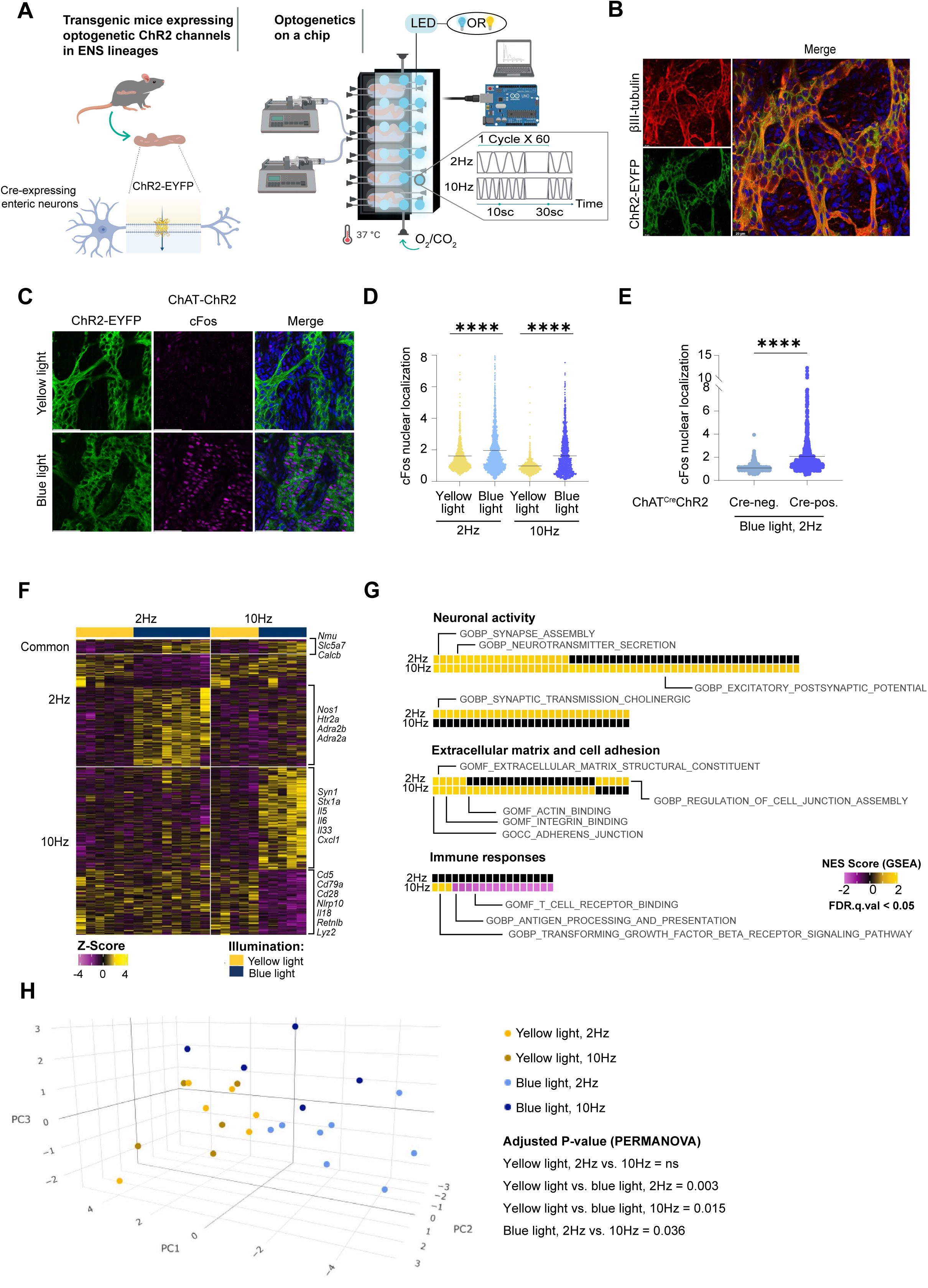
Optogenetic activation of enteric cholinergic neurons differentially modulate colonic transcription, in a frequency-dependent manner. **(A)** Schematic description of the optogenetics-integrated gut organ culture system. Intact intestinal tissues are dissected from mice expressing ChR2 in Cre-positive neurons, and connected to input and output ports of the gut culture device. Pumps controlling medium flow inside the lumen and in the external medium chamber. The lid incorporates blue or yellow LED light diodes and O2/CO2 input ports. LED illumination patterns in each channel are controlled by Arduino electronic platform and custom-made software. **(B)** Whole-mount staining of the colonic ENS in ChAT-ChR2 mice. Green: ChR2-EYFP; Red: βIII-tubulin. Scale bar: 20 µm. **(C-E)** Nuclear localization of cFos following optogenetic activation of ChAT-ChR2 neurons. (C) Representative whole-mount staining images of colonic myenteric plexus in cultured tissues illuminated by yellow or blue light (10Hz) and collected after 2h in culture. Magenta: cFos; Green: ChR2-EYFP. Scale bar: 50 µm. (D-E) Quantification of cFos nuclear localization following optogenetic stimulation of ChAT-ChR2 (homozygote) (D) or ChAT(Cre-positive/negative)-ChR2 (E). 50 images were analyzed, acquired from 13 gut tissues in 4 independent experiments. Statistical significance was determined by one-way ANOVA with multiple comparisons (D) or student’s t-test (E): ****p<0.0001. **(F)** Heatmap (z-score) of differentially expressed genes (Log2FC>=0.58, p<=0.05) in ChAT-ChR2 tissues illuminated by yellow or blue light. Light was delivered for 30min, in 60 cycles of (10sec of 1ms pulses delivered at either 2 or 10Hz, followed by a 20sec break). Tissues were collected after 2h in culture. Common: genes significantly up- or down-regulated in both conditions; 2/10Hz: genes significantly changed only in the designated condition. Selected genes of interest are highlighted. **(G)** Gene set enrichment analysis (GSEA) identified pathways significantly up- or down-regulated following optogenetic stimulation by blue light, delivered at 2/10Hz stimulation patterns (compared with tissues illuminated by yellow light as an internal control). Significant pathways were grouped into relevant biological categories. The normalized enrichment scores (NESs) and false discovery rate (FDR) q values (<0.05) are indicated. **(H)** 3D Principal component analysis (PCA) of ENS-related genes, in optogenetics-stimulated tissues. Statistical significance was determined by non-parametric permutational multivariate analysis of variance (PERMANOVA).

We next generated transgenic mice that expresses the light-sensitive ion channel channelrhodopsin-2 (ChR2) in specific subsets of the ENS. We first focused on enteric cholinergic neurons, that control ILC2 activity and type-2 immunity^3–6^. For this purpose, we crossed Ai32 mice (homozygous for a *loxP*-flanked STOP cassette followed by channelrhodopsin-2/EYFP fusion gene) with ChAT-IRES-Cre knock-in mice, to generate ChAT-ChR2 homozygotes transgenic mice that express the ChR2-EYFP fusion protein in enteric cholinergic (ChAT+) neurons. Whole-mount staining of colonic tissues demonstrated co-localization of ChR2-EYFP with the neuronal marker beta-3 tubulin in the ENS (**Fig.1B**). EYFP expression in enteric neurons was further validated by tissue fractionation (myenteric plexus, lamina propria and intestinal epithelium), dissociation and flow cytometry (within CD9^high^CD45^neg^Sca1^neg^ enteric muscularis myenteric plexus (MMP) neurons^11^) (**Fig.S2A-B**). ChR2 enables rapid activation of neurons by illumination with blue light (450-490nm). Indeed, live imaging of light-stimulated colon tissues connected to the gut culture device demonstrated intracellular calcium influx within myenteric neurons in response to blue light illumination, but not yellow light control (**Fig.S2C-F**).

We next analyzed colonic responses to optogenetic activation of enteric neurons, across a range of physiologically relevant frequencies. Electrophysiological recordings of colonic ENS activity performed in previous studies identified homeostatic innervation at 2Hz^15,16^. Firing frequencies were significantly elevated under inflammatory conditions (i.e. infection or experimental colitis models ^17–19^). Accordingly, ChAT-ChR2 colonic tissues were connected to the gut culture device^10,14^ and illuminated with cycles of blue light (460nm) for neuronal activation, or yellow light (590nm) as an internal control (**Fig.S1E**). Light was delivered for 30min, in 60 cycles of (10sec of 1ms pulses^20^ delivered at either 2 or 10Hz, followed by a 20sec break). After an additional 1.5h culture in the dark (total of 2h culture), tissues were collected and analyzed. Whole mount staining demonstrated increased nuclear localization of the neuronal transcription factor and activation marker cFos in the myenteric plexus of tissues illuminated by blue light, compared with tissues illuminated by yellow light (**Fig.1C-D**; **Fig.S3A**). To exclude the possibility of non-specific neuronal activation by blue light alone, ChAT-ChR2 heterozygote tissues were dissected from Cre-positive and Cre-negative littermates (that do and do not express ChR2, respectively), and connected to the gut culture device. Here again, blue light illumination significantly increased cFos nuclear localization in ChAT^Cre-Positive^-ChR2 tissues (compared with ChAT^Cre-Negative^-ChR2 tissues; **Fig.1E**, **Fig.S3B**). Thus, precise optogenetic stimulations of intestinal organ cultures trigger a measurable activation of enteric neurons.

We next aimed to determine whether on-chip neuronal activation modifies intestinal gene expression, and to establish whether intestinal transcriptome is differentially modulated by distinct neuronal activity patterns. Interestingly, optogenetic activation of cholinergic neurons rapidly induced colonic transcription, as determined by whole-tissue RNA sequencing (RNAseq) (**Fig.1F**) (Transcriptional responses induced by blue-light illumination were not induced in tissues illuminated by yellow light, compared with tissues cultured in darkness (**Fig.S3C**)). Comparison of 2/10Hz activation patterns revealed both frequency-dependent and frequency-independent transcriptional programs. For example, the cholinergic neuropeptide *Nmu* was induced by all activation patterns, in addition to other genes highly expressed by enteric cholinergic neurons^21^, such as the neuropeptide CGRP-β (*Calcb*), and the choline transporter *Slc5a7* (mediates acetylcholine synthesis) (**Fig.1F**). Pathway enrichment analysis revealed a significant induction in neuronal processes, including synaptic transmission and neurotransmitter secretion in both 2Hz and 10Hz (**Fig.1G**, **Fig.S3D-E**). In contrast, neuronal transcripts as the nitrergic inhibitory neuron marker *Nos1*, the serotonin receptor 2A *Htr2a*, and the adrenergic receptors alpha 2a and 2b (*Adra2a/b*) were induced only by 2Hz stimulation. Increased illumination frequency to 10Hz enhanced the expression of neuronal genes (i.e. *Syn1* and *Stx1a*) and pathways related to neural excitation and neurotransmitter release (**Fig.1G**, **Fig.S3D-E**). Principle component analysis (PCA) of ENS-related genes^22^ revealed that distinct 2/10Hz stimulation patters triggered discrete profiles of ENS transcription (compared with yellow light illumination controls; **Fig.1H**).

Remarkably, we identified frequency-dependent expression of immunological programs. 10Hz (but not 2Hz) stimulation of cholinergic enteric neurons induced ILC2 and type-2 immunity programs, including *Il5*, *Il6*, *Il33*, and *Cxcl1* while suppressing genes and pathways related to barrier defense (i.e. *Nlrp10*, *Il18*, *Retnlb*, *Lyz2*, *Cldn4*) lymphocytes activity (i.e. *Cd5*, *Cd79a*, *Cd28*) and antigen processing and presentation (**Fig.1F-G**, **Fig.S3E**). Thus, distinct firing patterns of enteric cholinergic excitatory motor neurons differentially control colonic neuro-immunological transcriptional programs.

To determine whether colonic transcription is differentially modulated by distinct neuronal ENS lineages, we examined tachykinin (*Tac1*)-expressing excitatory neurons (the *Tac1*-encoded neuropeptide Substance P controls Tregs development^10,11^ and regulates tissue protection^23^). We generated Tac1-ChR2 heterozygotes transgenic mice where expression of Cre recombinase results in transgene positive and negative littermates (that do and do not express ChR2, respectively). Tac1-ChR2 colonic organ cultures were illuminated by blue light (for 30min, in 60 cycles of (10sec of 1ms pulses^20^ delivered at either 2 or 10Hz, followed by a 20sec break)). Tissues were collected at 2h, and the transgene negative tissues served as an internal control (**Fig.2A**). cFos nuclear localization in the myenteric plexus was significantly increased in the Tac1^Cre-Pos^ChR2 transgene positive tissues (compared with Tac1^Cre-Neg^ChR2 tissues; **Fig.2B-C**, **Fig.S4A**). In contrast to cholinergic neurons, 10Hz optogenetic stimulation of Tac1-ChR2 neurons induced transcription of type-1 immune responses and barrier defense programs (**Fig. 2D-E**, **Fig.S4B**). These include *Ifng* (interferon-γ), the ENS-derived cytokine *Il18* (*Ifng*-inducing factor, controls barrier immunity^12,13^), the IL18-dependant anti-microbial peptide *Retnlb*^13,24^, and the antimicrobial peptides *Reg3b* and *Reg3g* (**Fig. 2D**). Additional epithelial barrier defense factors included the antimicrobial and IFNg-induced chemokine *Cxcl10*, the pathogenic Th17 cells inducer *Saa1* (serum amyloid A^25^), and the reactive oxygen species-producing enzymes *Nos2*, *Duox2* and *Duoxa2* (**Fig. 2D**). In agreement, pathways related to interferon signaling and defense response to symbiont were highly enriched (**Fig 2E**, **Fig.S4B**). Thus, distinct ENS lineages differentially modulate colonic expression of immunological programs.

**Figure 2:**
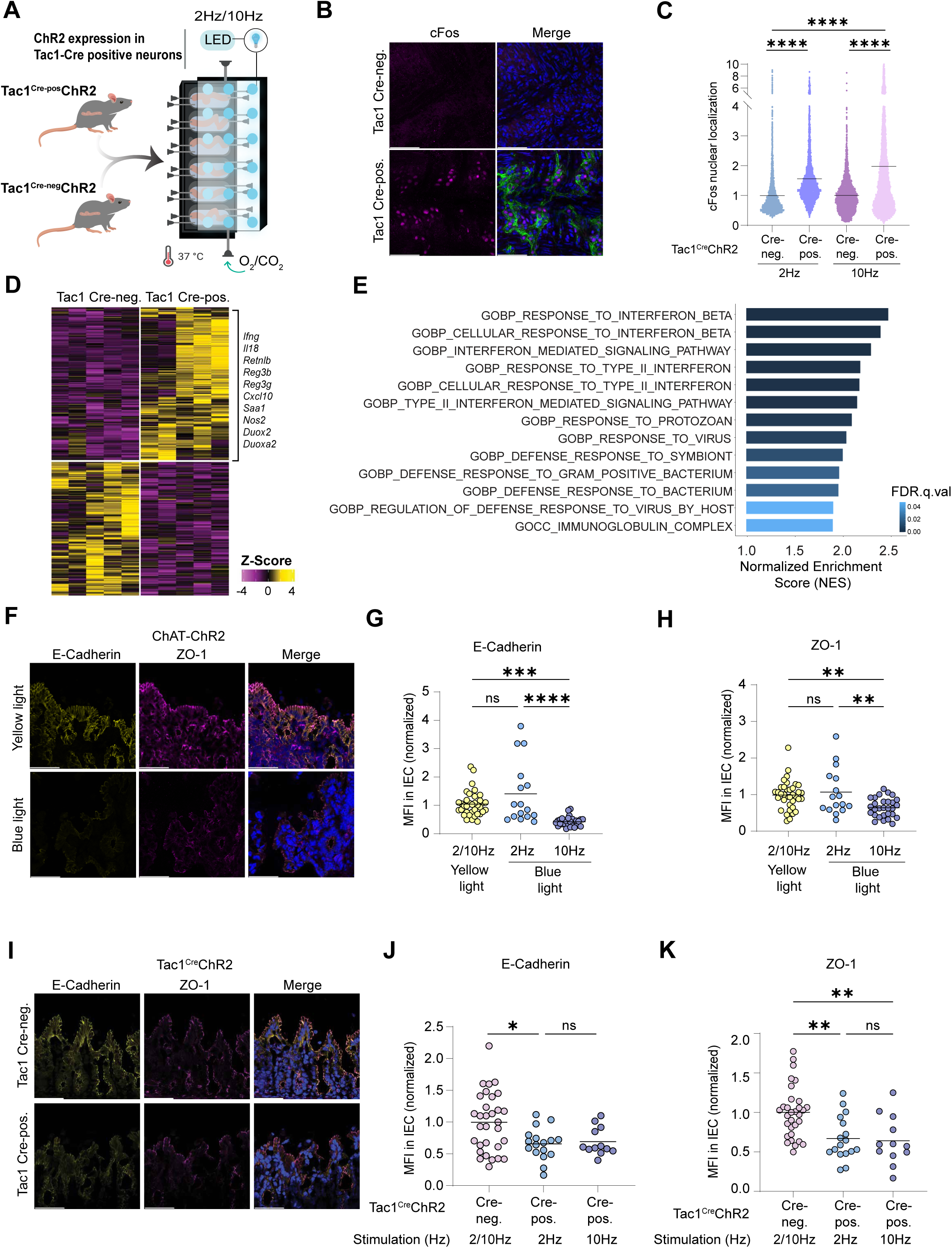
Optogenetic activation of enteric tachykinin-expressing neurons demonstrates lineage-dependent expression of immunological programs. **(A)** Experimental outline: Colon tissues collected from Tac1-Cre-transgene positive or negative mice and illuminated by blue light. Light was delivered for 30min, in 60 cycles of (10sec of 1ms pulses delivered at either 2 or 10Hz, followed by a 20sec break). Tissues were collected after 2h in culture. **(B-C)** Nuclear localization of cFos following optogenetic activation of Tac1-ChR2 neurons. (B) Representative whole-mount staining images of colonic myenteric plexus in cultured tissues illuminated by blue light (10Hz, per 2A). Magenta: cFos; Green: ChR2-EYFP. Scale bar: 50 µm. (C) Quantification of cFos nuclear localization following optogenetic stimulation. 69 images were analyzed, acquired from 14 gut tissues in 3 independent experiments. Statistical significance was determined by one-way ANOVA with multiple comparisons: ****p<0.0001. **(D)** Heatmap (z-score) of differentially expressed genes (Log2FC>=0.58, p<=0.05) in Tac1-Cre-transgene positive or negative mice tissues illuminated by blue light (10Hz, per 2A). Tissues were collected after 2h in culture. Selected genes of interest are highlighted. **(E)** Gene set enrichment analysis (GSEA) identified pathways significantly enriched in Tac1-Cre-positive tissues (compared with Tac1-Cre-negative). Top pathways from the M5 canonical ontology gene set database are shown. The normalized enrichment scores and false discovery rate (FDR) q values <=0.05; scale bar) are indicated. **(F-K)** Representative Confocal microscopy images of ChAT-ChR2 (F) and Tac1-Cre-ChR2 (I) tissues immunostained for epithelial markers. Yellow: E-cadherin; Magenta: ZO-1 (scale bar: 50µm). (J-H, J-K): Quantification of E-cadherin (G and J) and ZO-1 (H and K) mean fluorescence intensity (MFI) in the intestinal epithelial cells (IEC), in individual images.

Emerging evidence suggest that the ENS coordinates epithelial integrity and barrier protection^12,13,26^. To determine whether on-chip activation of distinct neuronal subtypes and activation patterns affects epithelial tight junction (TJ) integrity, homozygote ChAT-ChR2 and heterozygote Tac1-ChR2 colons were optogenetically activated at 2/10Hz. Interestingly, cholinergic activation of ChAT-ChR2 colons at 10Hz, but not 2Hz, rapidly compromised TJ integrity, as indicated by a significant decrease in epithelial ZO-1 and E-cadherin protein staining (**Fig. 2F-H**; **Fig.S3F**). In contrast, optogenetic activation of Tac1+ neurons suppressed TJ protein expression in both 2Hz and 10Hz frequencies (**Fig. 2I-K**), suggesting that distinct ENS lineages differentially control epithelial barrier integrity in an innervation frequency-dependent manner.

Under physiological conditions, intestinal immunocytes and enterocytes are simultaneously exposed to numerous microbial and neuronal-derived signals. Potentially, cellular integration of combinatorial signals may drive diverse synergistic or antagonistic outcomes. Yet intestinal responses to defined combinations of microbial and neuronal signals are mostly unknown and are extremely challenging to investigate using existing methodologies. In contrast, the optogenetics-integrated gut organ culture system facilitates precise luminal infusion of gut microbes in parallel to optogenetic-mediated ENS stimulation, thus accelerating combinatorial neuro-immune-microbiota research at the whole-tissue level.

We examined colonic responses to the human symbiont *Clostridium ramosum*, a potent Treg-inducing microbe^27^ which we have previously found to modulate ENS activity to promote anti-inflammatory immune responses^1,10^. Cultured ChAT-ChR2 colons were illuminated with blue or yellow light (10Hz, pattern described above), in parallel to luminal infusion of *C. ramosum* or *Peptostreptococcus magnus* (that does not induce Tregs development^27^) as an internal control (**Fig.3A**). Interestingly, despite vigorous optogenetic neuronal stimulation, Treg-inducing *C. ramosum* reduced cFos nuclear localization intensity in myenteric neurons (**Fig.3B-C**, **Fig.S5A**). Further, *C. ramosum* differentially modulated optogenetics-induced colonic transcription (**Fig.3D-E**). For example, optogenetics-mediated induction of cholinergic neuropeptides (e.g. *Nmu*, *Calcb,* and *Scg3*) and the suppression of barrier defense (i.e. *Nlrp10*, *Lyz2*, *Cldn4*) and lymphocytes activity (i.e. *Cd5*, *Cd79a*, *Cd28*) related genes were effectively eliminated in optogenetics-stimulated tissues infused with *C. ramosum* (**Fig.3D**). In contrast, luminal *C. ramosum* did not affect optogenetics-mediated induction of type-2 cytokines (as *Il5*, *Il6*, or *Cxcl1*) (**Fig.3D**). Yet, combinatorial optogenetic stimulation and luminal *C. ramosum* infusion remodeled neuro-immunological responses induced by optogenetic stimulation alone, mainly affecting genes and pathways related to cytokines production, binding, and cell-cell contact (**Fig.3D-E**, **Fig.S5B**). Finally, luminal infusion of *C. ramosum* (but not *P. magnus*) ameliorated epithelial TJ disruptions induced by 10Hz cholinergic innervation, suggesting a potential role for gut microbes in regulating enteric neuro-epithelial connections (**Fig.3F-G**, **Fig.S5D-E**).

**Figure 3:**
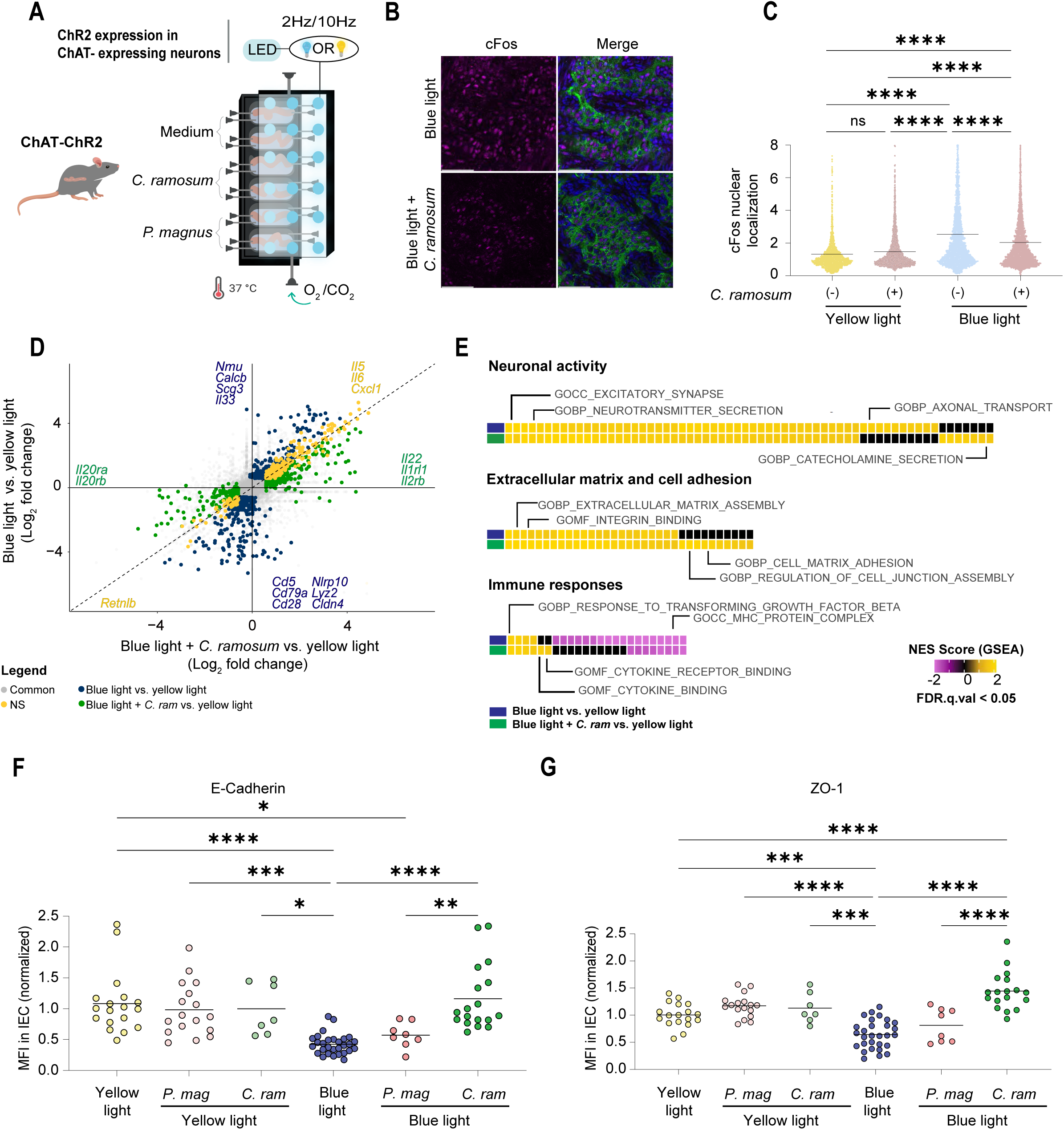
Luminal infusion of *C. ramosum* modulates optogenetic-induced transcriptional programs. **(A)** Experimental outline: Colon tissues collected from ChAT-ChR2 mice were infused with *C. ramosum*, *P. magnus*, or sterile medium, and illuminated by yellow or blue light. Light was delivered for 30min, in 60 cycles of (10sec of 1ms pulses delivered at either 2 or 10Hz, followed by a 20sec break). Tissues were collected after 2h in culture. **(B-C)** Nuclear localization of cFos following *C. ramosum* infusion and optogenetic activation of ChAT-ChR2 neurons. (B) Representative whole-mount staining images of colonic myenteric plexus in cultured tissues illuminated by blue light (10Hz, per 3A), with or without *C. ramosum*. Magenta: cFos; Green: ChR2-EYFP. Scale bar: 50 µm. (C) Quantification of cFos nuclear localization following bacterial infusion and optogenetic stimulation. 87 images were analyzed, acquired from 23 gut tissues in 3 independent experiments. Statistical significance was determined by one-way ANOVA with multiple comparisons. ****p<0.0001. **(D)** Comparison of transcriptional responses to cholinergic optogenetic stimulation (10Hz, per 3A), with or without luminal infusion of *C. ramosum* (relative to yellow light-illuminated control cultures). Common responses are labeled in yellow; responses unique to blue light illumination are labeled in blue; those unique to *C. ramosum* & blue light are labeled in green. Gene expression analysis of n=5 tissues (yellow light), n=5 tissues (blue light), n=6 tissues (blue light & *C. ramosum*) (Log2FC>=0.58, p<=0.05) is shown. Selected genes of interest are highlighted. **(E)** Gene set enrichment analysis (GSEA) identified pathways significantly up- or down-regulated following optogenetic stimulation by blue light, with or without luminal infusion of *C. ramosum* (relative to yellow light-illuminated control cultures). Significant pathways were grouped into relevant biological categories. The normalized enrichment scores (NESs) and false discovery rate (FDR) q values (<0.05) are indicated. **(F-G)** Quantification of Mean fluorescence intensity (MFI) of E-cadherin (F) and ZO-1 (G) in intestinal epithelial cells (IEC), in individual images (for representative image see in Figure S5). Statistical significance was determined by one-way ANOVA with multiple comparisons, adjusted p values: ****p < 0.0001; ***p < 0.001; **p < 0.01; *p < 0.05.

Taken together, our study introduces a unique experimental system enabling precise, real-time control of both enteric neuron activity and luminal microbiota composition at the whole-tissue level. We leverage this system to demonstrate that distinct neuronal lineages and their dynamic activity patterns differentially regulate intestinal gene expression and epithelial barrier integrity. Furthermore, we propose that combinatorial signals delivered by gut microbes and enteric neurons are locally integrated to regulate intestinal immunity and barrier defense.

The optogenetics-integrated gut organ culture system enables the systematic discovery and mechanistic exploration of novel neuroimmune and neuroepithelial connections. Furthermore, it facilitates the investigation of how these connections are modulated by drugs, microbes, or metabolites. This capability may open new avenues for streamlining drug discovery by enabling the rational selection and prioritization of candidate compounds identified through simple *in vitro* screens, prior to pre-clinical models.

## Methods

### Mice

Transgenic mice (based on C57BL/6 mouse background) were obtained from Jackson lab and maintained in the specific pathogen-free (SPF) mice facility at Bar-Ilan University (Israel). Mice were handled following protocols approved by the Bar-Ilan University ethics committee (ethics approval number BIU-BIU-IL-2203-131-1). Mice expressing channelrhodopsin-2/EYFP [RCL-ChR2(H134R)/EYFP] fusion protein in the Rosa26 locus downstream of a floxed-STOP cassette (Ai32 mice; RRID: IMSR_JAX:024109) were crossed with either ChAT-Cre (RRID: IMSR_JAX:006410) or Tac1-Cre (RRID: IMSR_JAX:021877) mice. Three lines of mice were generated: (1) ChAT-ChR2 homozygotes transgenic mice that express the ChR2-EYFP fusion protein in enteric cholinergic (ChAT+) neurons; (2) ChAT-ChR2 heterozygote mice in which Cre-positive and Cre-negative littermates do and do not express ChR2 (respectively); and (3) Tac1-ChR2 heterozygotes transgenic mice where expression of Cre recombinase results in transgene positive and negative littermates (that do and do not express ChR2, respectively). Tissues dissected from littermate mice were used in all experiments.

### Genotype characterization

Mice genotyping was performed according to the Jackson Laboratory website. Briefly, 1mm of mice tails were collected and dissolved at 55°C overnight in lysis buffer (100nM Tris-HCl Ph 8.5 (Fisher, 77-86-1), 5mM EDTA (Bio-Lab 009012230100), 0.2% SDS (Bio-Lab, 001981232300), 200nM NaCl (Millipore Cat#1064041000) dissolved in H2O) with 0.2mg/ml proteinase K (Bio-Lab, 1673238300). Then, deactivated at 65°C for 20 minutes and centrifuged for 10 minutes at max speed. 300µL of supernatant was collected to a new tube with an equal amount of isopropanol (J.T.Baker, 8067) and centrifuged at max speed for 8 minutes. The supernatant was discarded, DNA was resuspended in 100ul of T.E buffer and incubate for 1hr at 37°C. PCR program was executed according to strain-specific protocols found at the Jackson Laboratory website. Primers used:

ChR2-EYFP WT forward: 5¢-AAG GGA GCT GCA GTG GAG TA-3¢, reverse; 5¢-CCG AAA ATC TGT GGG AAG TC-3¢.

ChR2-EYFP mutant forward: 5¢-ACA TGG TCC TGC TGG AGT TC-3¢, reverse; 5¢-GGC ATT AAA GCA GCG TAT CC-3¢.

Tac1^Cre^ WT forward: 5¢-GCA TGT TTC CTG TTT CGT GA -3¢.

Tac1^Cre^ mutant forward: 5¢-TGG TGG CTG GAC CAA TGT-3¢.

Tac1^Cre^ mutant+WT reverse: 5¢-GCA TAT TTG GCT TTT ACT CTG G -3¢. ChAT^Cre^ WT forward: 5¢-GCA AAG AGA CCT CAT CTG TGG A -3¢.

ChAT^Cre^ mutant forward: 5¢-CAA AAG CGC TCT GAA GTT CCT -3¢. ChAT^Cre^ mutant+WT reverse: 5¢-CAG GGT TAG TAG GGG CTG AC -3¢,

### Optogenetics-integrated Gut Organ Culture System

The gut culture device was modified to support on-chip optogenetic stimulation (Supplementary Figure 1). Fabrication of the gut organ culture device and gut organ culture experiments (Yissachar et al., Cell, 2017) were performed with the addition of carbon black (Holland-Moran Cas#1333-86-4) into the PDMS (184 ®SYLGARD#761036) mixture, for fabrication of an opaque gut culture devices. We developed a three-layer culture device (Fig.S1A): (1) A standard organ culture device at the bottom, containing 6 intestinal organ cultures; (2) middle section adapter, for the flow of filtered and humidified medical-grade 95% O2 / 5% CO2 gas mixture; and (3) top section containing blue or yellow LED diodes controlled by Arduino UNO platform, to control LED illumination patterns (Fig.S1B). All experiment were conducted in the dark.

For gut culture experiments, intact whole colons were dissected sterilely from 14-day-old transgenic mouse littermates reared under SPF conditions. The solid lumen content was gently flushed, and the gut fragment was threaded and fixed over the luminal input and output ports of the gut organ culture device, using sterile surgical thread. The culture device was placed in a custom-made incubator that maintains a temperature of 37 °C, and tissue was maintained half-soaked in sterile, serum-free culture medium (Iscove’s Modified Dulbecco’s Medium without phenol-red (IMDM, GIBCO) supplemented with 20% KnockOut serum replacement (GIBCO), 2% B-27 and 1% of N-2 supplements (GIBCO), 1% L-glutamine, 1% non-essential amino acids, 1% HEPES) using a syringe pump. Tissues were infused with sterile culture medium without phenol-red into the gut lumen using a syringe pump, with or without purified bacterial cultures (*C. ramosum* or *P. magnus*, at 10^7–8^ CFU/mL). Gas outlet in the device lid enabled the flow of a humidified and filtered, medical grade 95% O2 / 5% CO2 gas mixture into the device. Experiments were terminated at 2h post-stimulation. After culture, tissues were subjected to further analysis (including RNAseq, immunofluorescence staining and imaging).

### Stimulation of optogenetic neurons using the gut culture device

Tissues were illuminated using blue or yellow LED diodes (5mm, 3.5V ∼ 4V) (Fig.S1E) connected to the Arduino UNO platform and placed above the gut organ cultures. Illumination pattern was controlled by custom-made software (Fig.S1C-D). Tissues were illuminated for 30 minutes, in 60 cycles of (10sec of 1ms pulses delivered at either 2 or 10Hz, followed by a 20sec break) (per Schiller et al., Immunity, 2021). Following optogenetic stimulation, tissues remained in culture for 1.5h, and collected at 2h post-stimulation.

### Bacterial cultures

*C. ramosum* and *P. magnus* were kindly provided by the Mathis/Bensoit lab at Harvard Medical School (Yissachar et al., Cell, 2017). The bacterial taxonomic classification was further validated by Sanger sequencing for the 16S gene. Microbes were cultured on Brucella Agar+5% Defibrinated Sheep Blood plate and incubated overnight (*C. ramosum*) or 48h (*P. magnus*) under anaerobic conditions at 37°C. Next, a single colony was isolated and culture in rich liquid medium (2% proteose peptone (OXOID Cat#LP0085), 0.5% NaCl (Millipore Cat#1064041000) and 0.5% yeast extract (ACROS, 451120010) supplemented with 250 mg/mL glucose (Millipore, 1083371000), 250 mg/mL K2HPO4 (Fisher scientific, 231-834-5), 50 mg/mL L-cysteine (SIGMA, C1276), 5 mg/mL Hemin (Millipore Cat#3741) and 5μl/ml vitamin *K*1 (Millipore, 5018900001)) in an anaerobic chamber. For organ culture experiments, bacterial cultures were resuspended for a final concentration of 10^7–8^ CFU/mL in sterile organ culture medium.

### Immunofluorescence staining and imaging

#### Whole mount staining of the myenteric plexus

Tissues were fixed in 4% paraformaldehyde (SIGMA, 252549) for 1 hour at 4°C, washed by PBSx1 (GIBCO, 14190-094) and permeabilized with 0.5% Triton X-100 for 2hr at 4°C. Blocking solution containing 10% donkey serum (Merck, D9663), 5% BSA (Millipore,821006), and 0.1% Triton X-100 (Millipore,1086031000) was added for 1hr at room temperature (RT). Primary antibodies anti-βIII-tubulin (Abcam ab41489), cFos (Cell Signaling Technology, 2250), and HuC/D (Santa Cruz SC-515624) were diluted in blocking solution (1:100) and incubated with tissues overnight at 4°C. The next day, tissues are washed 3 times with PBS-T (0.1% tween in PBSx1) and incubated overnight with secondary antibodies cy3-AffiniPure Donkey Anti-Rabbit IgG (Jackson lab, #711-165-152) and cy5 (Abcam, ab97147) diluted in blocking solution (1:200). After washing 3 times with PBS-T, tissues stained with DAPI (1:1000 in PBSx1, Mercury #1246530100) for 15 minutes at RT. Finally, tissues are mounted on slides with Fluoroshield mounting medium (SIGMA, F6182), and imaged using a confocal fluorescence microscope Leica Stellaris 5 (Leica Microsystems).

#### Immunofluorescence staining of tissue sections

For tissue section staining, intestinal tissues were frozen in OCT (labotal, CAT-14020108926) at -80°C. 7μm-thick sections of fresh-frozen tissues were sliced using a cryostat (CM1950). The sections were fixed with 100% cold acetone for 20 minutes, washed with PBSx1, and blocked (10% donkey serum, 0.1% Triton in PBSx1) for 1h at room temperature. Primary antibodies for ZO-1 (Invitrogen cat: 40–2200) 1:100 and E-cadherin (BioLegend cat:147302) 1:200, were diluted in blocking solution and incubated overnight at 4°C. The next day, slides are washed 3 times with PBS-T (0.1% tween in PBSx1) and incubated with a secondary antibody Cy5 (Jackson cat: 712-175-153) 1:250; Cy3 (Jackson cat: 711-165-152) 1:500 diluted in PBS-T (0.1% tween) for 1h at room temperature. After washing 3 times with PBS-T, tissues were stained with DAPI (1:1000 in PBSx1) for 15 minutes at RT. Finally, tissues are mounted on slides with a Fluoroshield mounting medium (SIGMA, F6182). Tissue sections were visualized using a confocal fluorescence microscope Leica Stellaris 5 (Leica Microsystems), processed and analyzed using ImageJ software.

### Image analysis

Quantification of cFos nuclear localization in whole mount images was performed using custom ImageJ macros, tailored for ChAT-ChR2 or Tac1-ChR2 tissues, respectively.

The ChAT-ChR2 tissue macro employs a multi-step process. First, the DAPI channel was segmented, after being smoothed via Gaussian blur (sigma=2), and touching nuclei were separated with the watershed function. YFP channel (expressed in cholinergic enteric neurons) segmentation was done after applying the smoothing (gaussian blur; sigma=5) and fill holes functions. In order to verify the segmentation of the nuclei were eroded the DAPI mask by 1. Then, YFP positive nuclei were identified by using image Calculator function (And) to keep only overlapping areas of both binary masks. In order to measure the cytoplasmic cFos MFI, the nuclei outlines were enlarged, a new binary mask generated, with the original DAPI mask being subtracted via image calculator function (subtract). Finally, the cFos MFI was quantified for both the nuclear and cytoplasmic regions by applying the analyze Particles function on the cFos channel. The ratio of nuclear to cytoplasmic MFI was presented to account for variations in cFos expression between cells.

The Tac1-ChR2 tissue macro works in a similar fashion. Following manual verification of proper spatial localization in the tissue, via presence of HuC/D staining, the macro quantifies the cFos MFI within all nuclei present in each field of view.

In both analyses, the measured cFos MFI is normalized to the mean of the control MFI values to account for potential inter-experimental variations in staining intensity. Additionally, manual verification of identified nuclei was conducted to minimize inclusion of false positives.

### Flow cytometry

#### Colon dissociation

The large intestine was fractionated into three different layers: ENS, lamina propria, and intestinal epithelial cells, as detailed below:

#### ENS isolation

Colons were opened lengthwise, and the muscularis myenteric plexus was dissected completely under a binocular microscope using watchmaker forceps (Zhang and Hu, 2013) and placed in a 1.5mL tube containing 0.1mg/mL Liberase TL (54101135001, Sigma) in neutral Hank’s Balanced Salt Solution (HBSS, 02-018-1A, Biological Industries) to digest for 0.5 h at 37°C, and then mechanically disrupted by mild shaking. Tissue was then further digested with 0.05% Trypsin -EDTA solution at 37°C for 10 min. After trituration using a P1000 pipette, single cell suspensions were filtered through 70um filter, cleaned of debris by centrifugation through 1ml fetal bovine serum (110g for 10min). This cells were then centrifuged and counted, for staining and flow cytometry.

#### IEC and LP isolation

Intestinal pieces incubate in fresh, pre-warmed extraction medium (5% DTT (fisher, BP172-5), 0.5M EDTA (Bio-Lab 009012230100), FBS (SIGMA, F9665) and RPMI (SIGMA, R7509)) for 10min at 37°C in a urine cup with a stir bar (per tissue, stir at 500 rpm). The extraction medium containing the IEC layer was collected and stained for flow cytometry. For isolation of LP lymphocytes, the intestinal pieces were washed in RPMI (37°C) and rolled over paper towels to remove mucus. Tissue was cut to small pieces using scissors and incubated in pre-warm digestion medium (FBS, Dispase (gibco cat. no 1705-041), Collagenase (Gibco cat. no 1701-015) and RPMI) for 30 min at 37°C (per tissue, stir at 500 rpm). The medium is filtered using a 40µm cell strainer, into a 50ml tube with 2ml FBS. Then rinse with an additional 20-25 ml of cold RPMI and 1 ml of FBS. Following centrifugation at 1500rpm for 10 min the cellular pellet was resuspended in 1ml, and filtered through a 40 µm cell strainer. This solution is then centrifuged and counted, for staining and flow cytometry.

#### Extracellular staining and flow cytometry

Purified single-cells suspensions were stained using antibodies against mouse CD45 (Miltenyi Biotec; 130-117-498), SCA1 (Miltenyi Biotec; 130-102-832) and CD9 (Miltenyi Biotec; 130-102-612) for 15min at 4°C. Cells were washed using RPMI and fixed (1% formalin in DMEM, overnight at 4°C). Cells were analyzed by BD LSRFortessa flow cytometer and data was processed with FlowJo software.

### Calcium Imaging

To monitor enteric neuronal activation through calcium signaling, we utilized gut tissue from a mouse expressing light-gated cation channel ChR2 under the ChAT promoter. The tissue was stained with the synthetic calcium indicator Rhod-2 (50 µg). Calcium Indicator Preparation: A solution containing 8 µl DMSO, 2 µl Pluronic acid, and 90 µl physiological saline was added to the vial of Rhod-2 and stirred for 30 minutes. Imaging Setup: The tissue was mounted on a microscope stage (Slicescope 6000, Scientifica) equipped with a CCD camera (EXI-Blue, QIMAGING). Perfusion with Ringer solution (composition specified) bubbled with 95% O2/5% CO2 was maintained throughout the experiment. Following a 10-minute adaptation period, the prepared calcium indicator solution was applied to the tissue surface at a final concentration of 2.5 µM. Perfusion was stopped for 15 minutes to facilitate efficient staining, then resumed. Light Stimulation and Data Acquisition: Light stimulation was used to induce neuronal activity. The resulting changes in fluorescence intensity were captured at a frame rate of 100 milliseconds, with 1-minute movies acquired for offline analysis. Data Analysis: Changes in fluorescence intensity were quantified by subtracting each acquired frame from the average of baseline frames (recorded during no stimulation). This difference was then normalized by dividing by the average baseline value. To assess enteric neuronal activity, the average fluorescence change within a defined region of interest (ROI) was calculated. It is important to note that this analysis might also capture induced muscle contractions, as they can also cause changes in pixel intensity.

### RNA sequencing and bioinformatics analysis

Colon tissue fragments (3 mm, proximal colon) were stored in RNAlater (Qiagen) overnight at 4c, and homogenized using a bead beater; then bulk RNA was extracted using RNeasy Plus Universal Mini QIAGEN kit. Libraries were prepared using a modified SMART-Seq2 protocol. In brief, 1ng of purified RNA was taken for reverse transcription with Maxima Reverse Transcriptase (Life Technologies) and whole-transcription amplification (WTA) with KAPA HotStart HIFI 2 × ReadyMix (Kapa Biosystems) for 20 cycles. WTA products were purified with Ampure XP beads (Beckman Coulter), quantified with Qubit dsDNA HS Assay Kit (ThermoFisher) and assessed with a high-sensitivity DNA chip (Agilent). RNA-seq libraries were constructed from purified WTA products using Nextera XT DNA Library Preperation Kit (Illumina). In order to increase the depth of sequencing, paired end reads were sequenced on 4 lane(s) of an Illumina NextSeq 500/550 three times by The Crown Genomics institute of the Nancy and Stephen Grand Israel National Center for Personalized Medicine,Weizmann Institute of Science.

Poly-A/T stretches and Illumina adapters were trimmed from the reads using cutadapt (Martin M., *EMBnet*, 2011) (version 2.7); resulting reads shorter than 30bp were discarded. Reads were mapped to the M. musculus reference genome GRCm39 using STAR (Dobin et al., *Bioinformatics*, 2013) (version 2.7.3a), supplied with gene annotations downloaded from Ensembl (https://www.ensembl.org/index.html) (and with EndToEnd option and outFilterMismatchNoverLmax was set to 0.04). Reads with the same UMI were removed using the PICARD MarkDuplicate tool using the BARCODE_TAG parameter. Expression levels for each gene were quantified using htseq-count (http://www-huber.embl.de/users/anders/HTSeq/doc/overview.html) (version 0.11.2), using the gtf above. Pipeline was run using snakemake (Koster et al., *Bioinformatics*, 2012). The output was ∼4 million reads per sample per Illumina NextSeq run. All three counts tables from the 3 independent sequencing runs were merged together, for a final sequencing depth of ∼10-12 million reads per sample. The initial processing described above done by Avital Sarusi-Portuguez from The Mantoux Bioinformatics institute of the Nancy and Stephen Grand Israel National Center for Personalized Medicine, Weizmann Institute of Science.

All gene reads were converted from Ensembl ID to gene symbol (in case of multiple Ensembl ID that were assigned to the same gene, the sum of total counts were calculated and assigned to the corresponding gene symbol). Differential gene expression analysis was performed using DESeq2 (Love M et al., *Genome Biology*, 2014) (1.36.0) R/bioconductor package. DESeq2 applies the Wald’s test on normalized counts and uses a negative binomial generalized linear model, which determines differentially expressed genes and log-fold changes. Significant differentially expressed genes were selected using threshold values of p-value <= 0.05 and abs(log2foldchange) >= 0.58 (Fold Change (FC)>=1.5). Heatmaps were generated for data visualization using ComplexHeatmap (Gu Z. et al., *Bioinformatics*, 2016) (version 2.12.1) R package. Briefly, variance stabilization transformation was applied, normalized expression for each gene in each sample was normalized to the average of the control group, and z-score was calculated and color-coded for generating the heatmaps. For comparison of ENS-expressed genes within optogenetic-activated tissues, genes within enteric neuron classes (per Morarach et al., *Nature Neuroscience*, 2021) that exhibited significant up- or down-regulation in response to optogenetic stimulation, were identified and extracted. Principal component analysis was done by using stats (version 4.2.1) R package, and 3-D PCA plot was generated using plotly (Sievert C*., Interactive Web-Based Data Visualization with R, plotly, and shiny*, 2020) (version 4.10.1) R package. Statistical analysis of permutational multivariate of variance was performed using pairwiseAdonis (Martinez Arbizu, P., 2020) package in R with pairwise.adonis() function with default parameters. Scatter plot was generated using ggplot2 (H. Wickham, *Elegant Graphics for Data Analysis. Springer-Verlag New York*, 2016) (version 3.4.2) R package. For pathway enrichment analysis, differentially expressed genes were analysed using Metascape (Zhou Y. et al., *Nat Commun,* 2019). Additionally, Gene Set Enrichment Analysis (Subramanian, Tamayo, et al., *PNAS*, 2005 and Mootha, Lindgren, et al., *Nature Genetics*, 2003) (GSEA version 4.3.2) was performed for all the genes ranked (-log10(p-value) /sign(log2FoldChange)), with M5 ontology gene sets (v2023.1). Relevant pathways from GSEA results with q-value (FDR)<=0.05 were plotted using ggplot2 or complexHeatmap. Non-significant pathways (FDR q>0.05) were assigned as NES=0. To evaluate the potential effect of yellow light illumination on colonic gene expression, we performed RNASeq of whole tissue samples following yellow light illumination at 10Hz compared with tissue that were not illuminated at all (darkness). Samples (n=3 per conditions) were used for library construction (using TruSeq stranded mRNA library prep kit) and sequenced at the Bar-Ilan University Scientific Equipment Center using the Illumina NextSeq platform (NextSeq 500 High Output v2 kit (#FC-404-2005)). Differential gene expression analysis was performed using DESeq2 as described above, and volcano plot was generated using ggplot2.

## Legends

**Supplementary figure 1:**
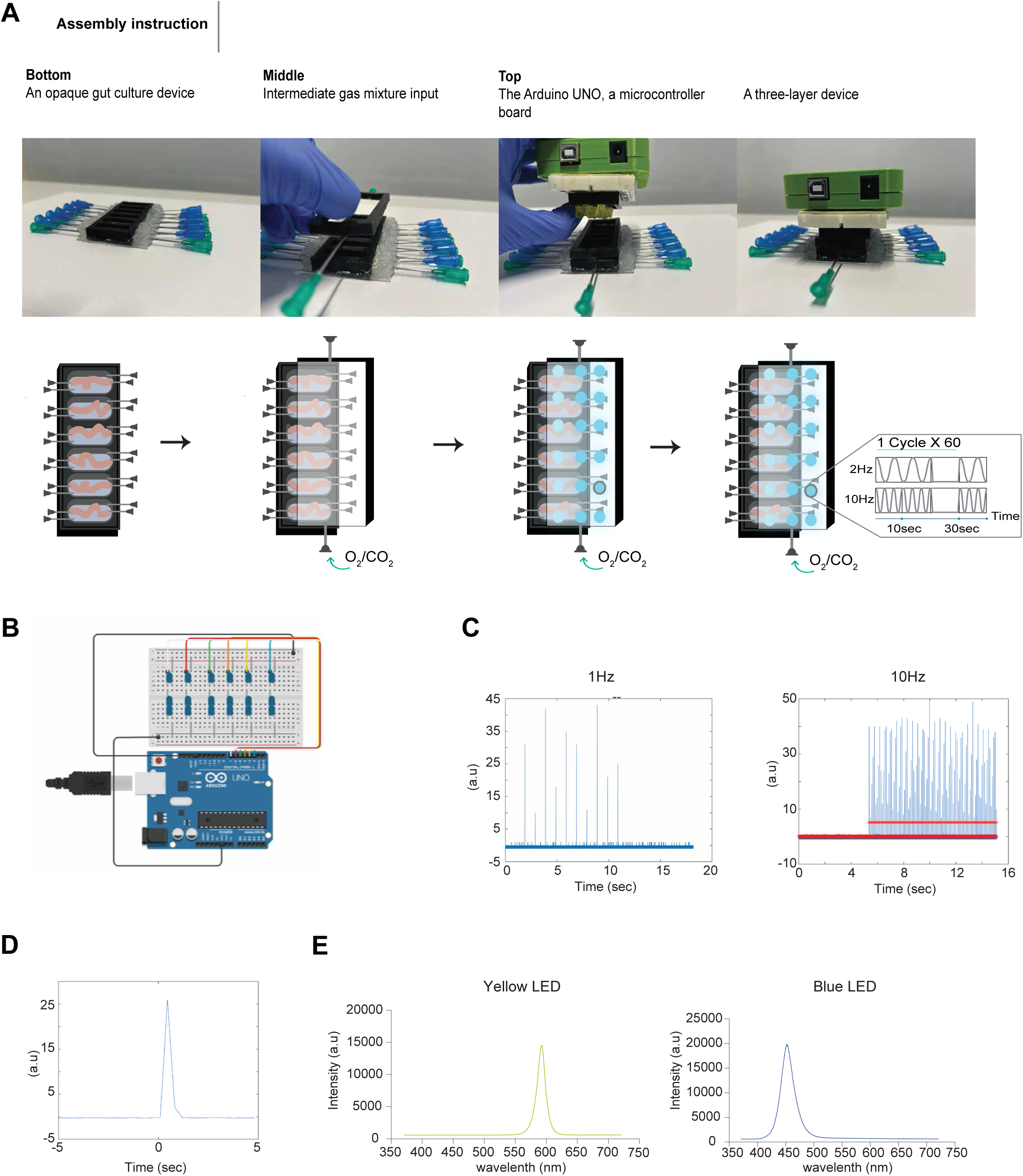
**(A)** Assembly instructions for three-layered, optogenetics-integrated gut organ culture device. **(B)** Blueprint of the LED array design. The blue and yellow LED light arrays were powered and controlled by an Arduino uno circuit, and custom-made software (supplementary data). **(C-D)** Illumination system validation. (C) Raw data with the detected pulses. Pulses were administered at 1 and 10Hz. (D) average pulse width (1msec). **(E)** Wavelength validation of both LEDs used, revealing the expected peaks of 570nm and 450nm for yellow and blue LEDs, respectively.

**Supplementary figure 2:**
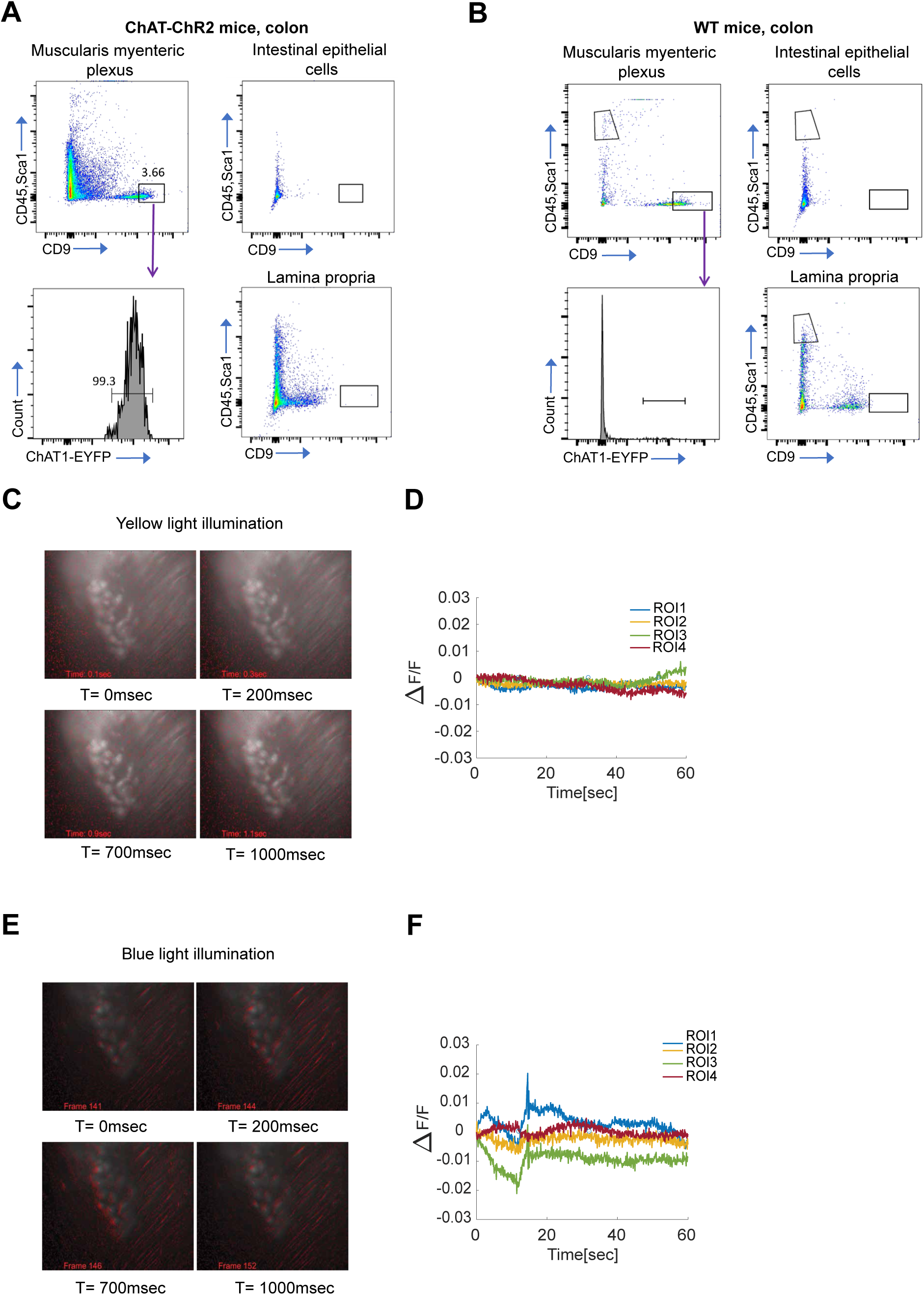
**(A-B)** Flow cytometry analysis of colonic epithelial layer, lamina propria, and muscularis myenteric plexus, dissected from ChAT-ChR2 (A) or wild-type (WT) (B) mice. Cells were stained using anti-CD45, SCA1 and CD9. ChAT-EYFP-expressing enteric neurons (CD45,Sca1^negative^, CD9^high^) are shown in the muscularis myenteric plexus of ChAT-ChR2 mice (bottom left). **(C and E)** Time lapse frame sequence following a train of 100msec pulses applied for 10 secs (stimulus onset is defined as t=0) using a set of LEDs, revealing the robust increase in calcium induced by the ChR2 following blue (F) but not yellow light (C) illumination, as well as the readily observed contraction of the tissue. **(D and F)** The average fluorescence signal change in the selected fields of view within the tissue, revealing the robust increase in calcium fluorescence signal following blue light illumination.

**Supplementary figure 3:**
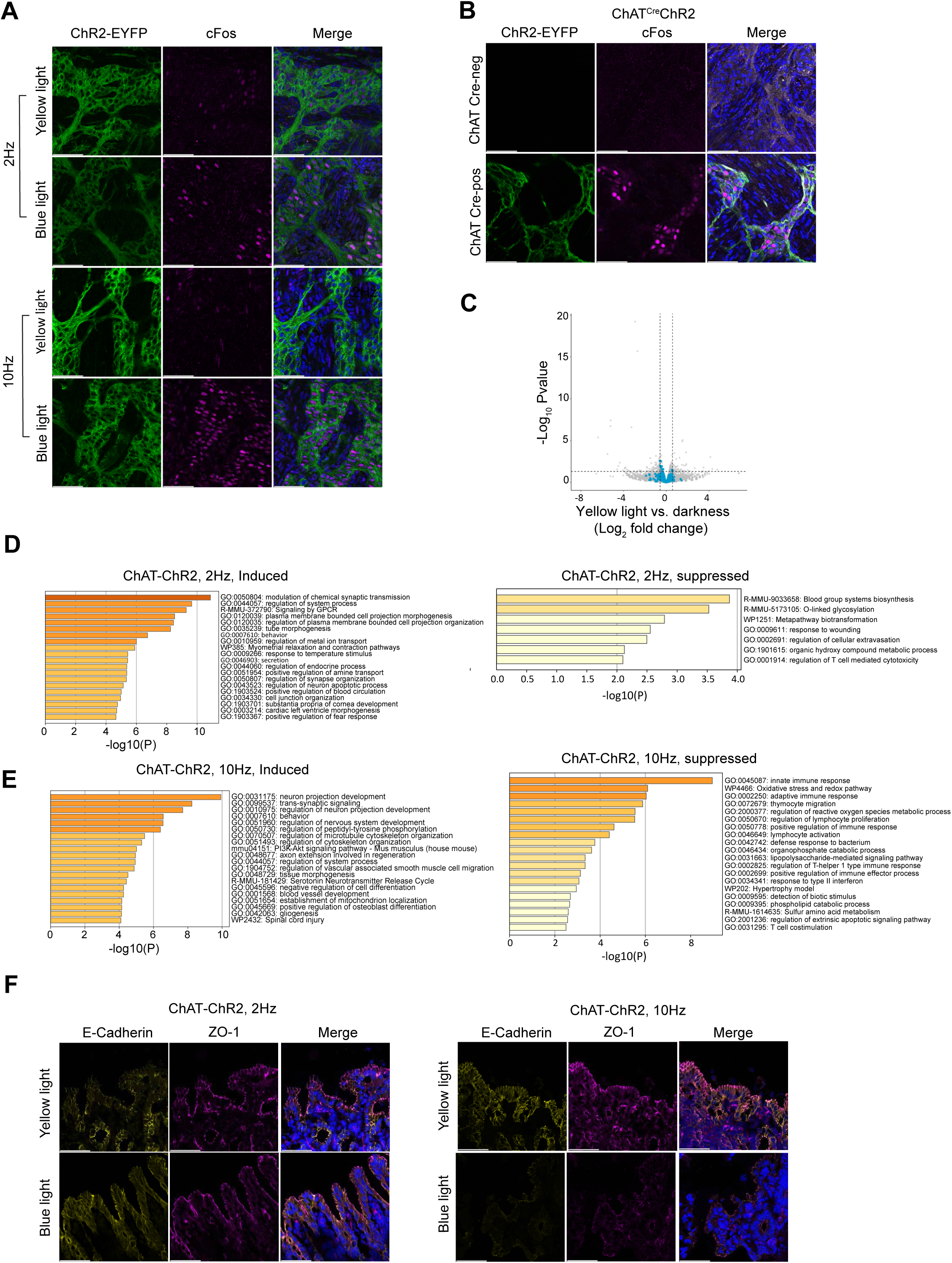
**(A-B)** Representative whole-mount staining images of colonic myenteric plexus in cultured tissues following optogenetic activation of ChAT-ChR2 (homozygote) (A) or ChAT (Cre-positive/negative)-ChR2 (B) enteric neurons. Magenta: cFos; Green: ChR2-EYFP. Scale bar: 50µm. **(C)** Volcano plot representation of gene expression comparing colon tissues from ChAT-ChR2 mice, cultured using the optogenetic-integrated gut culture system and illuminated by 10Hz yellow light versus darkness, for 2h. For comparison, significantly up-regulated genes following blue light illumination at 10Hz (per Fig.1F) are highlighted in blue. **(D)** Metascape pathway enrichment analysis of transcripts induced or suppressed in ChAT-ChR2 tissues illuminated by blue vs. yellow light, at 2 Hz for 2 hours. **(E)** Metascape pathway enrichment analysis of transcripts induced or suppressed in ChAT-ChR2 tissues illuminated by blue vs. yellow light, at 10 Hz for 2 hours. **(F)** Representative Confocal microscopy images of ChAT-ChR2 mice tissues illuminated by yellow or blue light (2/10Hz) immunostained for epithelial markers, Yellow: E-cadherin; Magenta: ZO-1 (scale bar: 50µm).

**Supplementary figure 4:**
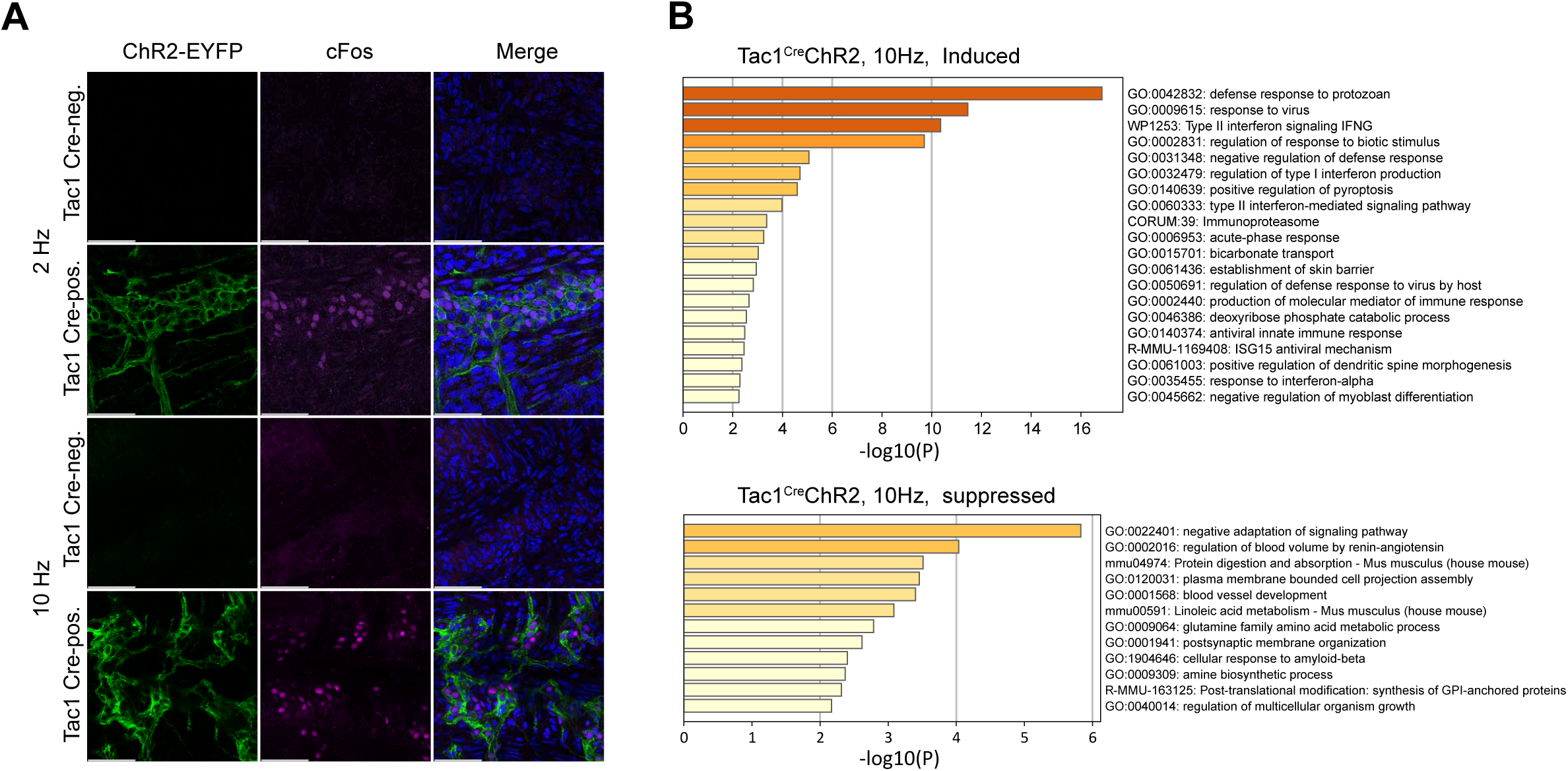
**(A)** Nuclear localization of cFos following optogenetic activation of Tac1-ChR2 neurons. Representative whole-mount staining images of colonic myenteric plexus in cultured tissues illuminated by blue light (2/10Hz) Magenta: cFos; Green: ChR2-EYFP. Scale bar: 50 µm. **(B)** Metascape pathway enrichment analysis of transcripts induced or suppressed in Tac1-Cre-positive tissues (compared with Tac1-Cre-negative) illuminated by blue light (10Hz).

**Supplementary figure 5:**
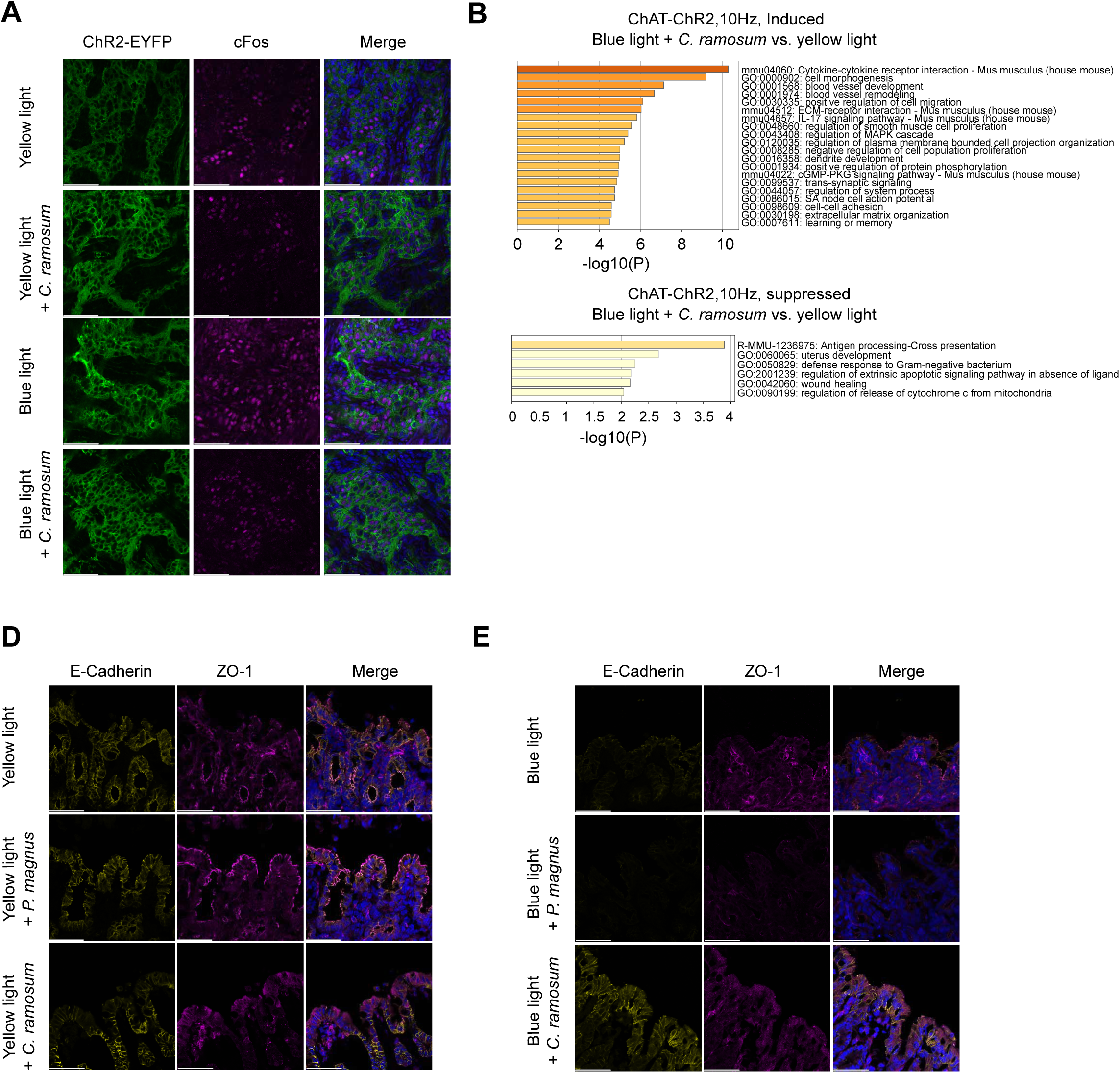
**(A)** Representative whole-mount staining images of colonic myenteric plexus in cultured tissues following optogenetic activation of ChAT-ChR2 tissues, with or without luminal infusion of *C. ramosum*. Magenta: cFos; Green: ChR2-EYFP. Scale bar: 50µm. **(B)** Metascape pathway enrichment analysis of transcripts induced or suppressed in ChAT-ChR2 tissues illuminated by blue light and infused with *C. ramosum* (compared with yellow light-illuminated tissues), at 10 Hz for 2 hours. **(D-E)** Representative Confocal microscopy images of ChAT-ChR2 mice tissues illuminated by yellow or blue light (10Hz) and infused with *C. ramosum* or *P. magnus*. Tissues immunostained for epithelial markers, Yellow: E-cadherin; Magenta: ZO-1 (scale bar: 50µm).

